# Assessment of endocytic traffic and Ocrl function in the developing zebrafish neuroepithelium

**DOI:** 10.1101/2022.06.14.496217

**Authors:** Daniel M. Williams, Lale Gungordu, Anthony Jackson-Crawford, Martin Lowe

**Affiliations:** School of Biological Sciences, Faculty of Biology, Medicine and Health, University of Manchester, The Michael Smith Building, Oxford Road, Manchester, M13 9PT, UK; School of Biosciences, University of Sheffield, Firth Court, Western Bank, Sheffield, S10 2TN, UK; European Research Institute for the Biology of Ageing, University Medical Centre Groningen, University of Groningen, Groningen, the Netherlands; Department of Blood Sciences, Grange University Hospital, Llanyravon, Gwent, NP44 8YN, UK

**Keywords:** endocytosis, neuroepithelium, OCRL, Lowe syndrome, megalin, zebrafish

## Abstract

Endocytosis is a vital process, required during development and for maintenance of tissue homeostasis, that allows cells to internalize a wide range of molecules from their environment as well maintain their plasma membrane composition. The ability to visualise endocytosis in vivo requires suitable assays to monitor the process. Here, we describe imaging-based assays to visualize endocytosis in the neuroepithelium of living zebrafish embryos. These assays rely on injection of fluorescent tracers into the brain ventricles followed by live imaging and can be used to study fluid-phase or receptor-mediated endocytosis, for which we use receptor-associated protein (RAP) as a ligand for LDL receptor-related protein (LRP) receptors expressed at the neuroepithelium. Using dual colour imaging combined with transient or stable expression of endocytic markers, it is possible to track the progression of endocytosed tracers and to monitor trafficking dynamics. Using these assays, we reveal a role for the Lowe syndrome protein Ocrl in endocytic trafficking within the neuroepithelium. We also find that the RAP binding receptor Lrp2 appears to only partially contribute to neuroepithelial RAP endocytosis. Altogether, our results provide a basis to track endocytosis within the neuroepithelium in vivo, and support a role for Ocrl in this process.

**Summary statement:** We describe live imaging assays to analyse endocytosis in the zebrafish neuroepithelium and show involvement of the inositol phosphatase OCRL in this process

## Introduction

Neuroepithelial cells and related cell types such as radial glia are polarised progenitor cells that make up the bulk of brain tissue at the early stages of brain development (Gotz and Huttner, 2005; Taverna et al., 2014). These cell types respond to multiple developmental cues that guide both tissue patterning and the post-mitotic fate neuronal progenitors commit to in order to establish a functional organ (Gotz and Huttner, 2005; Taverna et al., 2014). Multiple studies have demonstrated the importance of post-Golgi endocytic trafficking in the regulation of neuronal progenitor cell function and fate. Sensing of signals that influence neuroepithelial cell behaviour is in part mediated by receptors present at the apical surface, with cell surface receptor levels often maintained by receptor recycling from endosomes. As an example, apical enrichment of the recycling receptor LRP2 (Nagai et al., 2003) in mammalian neuroepithelial cells shapes tissue patterning within the forebrain through endocytosis of the secreted morphogen Sonic Hedgehog (SHH) with the absence of LRP2 leading to an impairment of SHH signalling during early neurogenesis in mice (Christ et al., 2012; Willnow and Christ, 2017). Bound ligands internalised into endosomes from the neuroepithelial cell surface can also influence neuroepithelial cell behaviour. The asymmetric segregation of the Notch ligand DeltaD within Rab5 positive SARA endosomes has been shown to be a key neuroepithelial cell fate determinant (Coumailleau et al., 2009; Kressmann et al., 2015; Nerli et al., 2020; Zhao et al., 2021) whilst the positioning of Rab11 recycling endosomes can control developmental signalling within the zebrafish retinal neuroepithelium and Drosophila sensory organ precursor cells (Clark et al., 2020; Emery and Knoblich, 2006; Furthauer and Gonzalez-Gaitan, 2009). Underscoring the importance of understanding how post-Golgi trafficking events regulate neuroepithelial cell function are mutations in multiple trafficking regulators that are linked to CNS abnormalities in mammals. Mutations in α-SNAP and ARF-GEF2 lead to an impairment of neuronal development through the over or underproduction of progenitor and neuronal cells respectively (Chae et al., 2004; Sheen et al., 2004), whilst mutations in enzymes that regulate phosphoinositide signalling on membranes, such as OCRL and INPP5E, lead to a variety of CNS associated symptoms in humans that include epilepsy, developmental delay and intellectual disability (Bokenkamp and Ludwig, 2016; Mehta et al., 2014; Parisi, 2009; Ramirez et al., 2012).

The direct study of endocytosis and downstream trafficking steps in neuroepithelial cells *in vivo* has been limited by the lack of established protocols allowing the application of many classical methods of studying endocytosis and endocytic trafficking used in cell culture. In zebrafish, the developing brain and its associated ventricular system are easily accessible and can be imaged by microscopy (Fame et al., 2020). In addition, transgenic zebrafish lines to examine the neuroepithelial endocytic pathway have been established and characterised, whilst additional studies have demonstrated the accessibility of the hindbrain ventricle for injection of fluorescent dyes (Clark et al., 2011; Fame et al., 2020; Lowery and Sive, 2005). Here, taking advantage of these features in zebrafish, we set out to establish a semi-quantitative endocytic uptake assay to investigate endocytosis and endocytic trafficking within the developing neuroepithelium. We use a fluorescently labelled high-affinity Lrp2 ligand (RAP) (Herz et al., 1991; Willnow et al., 1992) to assess both receptor-mediated endocytosis and downstream endocytic traffic in this tissue, and apply the assay to monitor endocytosis in *ocrl* and *lrp2* mutant zebrafish lines. We find that RAP uptake and downstream trafficking are defective in the *ocrl* mutant neuroepithelium, with loss of Ocrl also impacting on endosome size and levels of Lrp2 at the apical surface. These data suggest that the absence of Ocrl alters cell surface levels of endocytic receptors and that disruption of receptor trafficking may contribute to the neurodevelopmental defects seen in Lowe syndrome patients. Intriguingly, neuroepithelial RAP-uptake still occurs in *lrp2* null embryos suggesting that multiple RAP-binding receptors are present at the apical surface of neuroepithelial cells. Collectively, our results illustrate that the embryonic zebrafish neuroepithelium provides an ideal system in which to visualise and study endocytic trafficking *in vivo* in the context of neuronal development.

## Results

### Zebrafish neuroepithelial tissue endocytoses fluid-phase tracers injected into the hindbrain ventricle

Zebrafish brain development begins at the early stages of embryogenesis with the rudiments of the CNS present by 10 hours post-fertilisation (hpf) (Kimmel et al., 1995). By 24 hpf, the overall architecture of the fore, mid and hindbrain regions of the brain is well defined with tissue in these regions lining the fore, mid and hindbrain ventricles. At this stage of development, a significant proportion of brain tissue consists of neuroepithelial cells, a key progenitor cell type from which new-born neurons and further neuroepithelial cells are derived (Gotz and Huttner, 2005; Taverna et al., 2014). Viewed from both coronal (**Fig. 1A**) and sagittal (**Fig. 1B**) perspectives at 28 hpf, interkinetic nuclear migration of neuroepithelial nuclei along the apicobasal axis of the neuroepithelium gives the impression of a stratified tissue with multiple layers (Lee and Norden, 2013; Norden, 2017). Taking cells at the zebrafish midbrain hindbrain boundary (MHB) as a representative example however, neuroepithelial cells form a monolayer, adopting a narrow spindle like morphology that in fact spans the whole width of the brain, with the basal and apical processes of these cells contacting opposing surfaces. The basal process of neuroepithelial cells contacts the basal lamina adjacent to the brain wall, whilst the apical process forms junctions with neighbouring cells to integrate into an epithelial sheet that collectively makes contact with the cerebrospinal fluid (CSF) of the brain ventricles (**Fig. 1A,B**) (Fame et al., 2020; Gotz and Huttner, 2005; Norden, 2017; Taverna et al., 2014). Within these cells, the arrangement of endocytic compartments mirrors that seen in mammalian epithelial cell types. Early endosomal compartments are enriched towards the apical pole, whilst late endosomal compartments are more dispersed throughout the cell body, as previously described (Clark et al., 2011), and visualised live here using EGFP-tagged Rab5c and Rab7, respectively (**Fig. S1, see also Fig. 1D**). The early and late endosomes in the zebrafish neuroepithelium are dynamic, with Rab5c-positive early endosomes tracking from the periphery towards the cell interior, albeit with some bidirectional movement observed, and Rab7-positive endosomes displaying a more striking bidirectional movement deeper within the cytoplasm (**Movie 1 and 2**). We also visualized the vesicle coat protein clathrin, by expressing EGFP-tagged clathrin light chain (CLC). This revealed numerous small puncta close to or at the apical membrane and small puncta moving within the cytoplasm that likely correspond to endocytic pits and vesicles of the neuroepithelial cells (**Movie 3**). There were also larger dynamic puncta likely corresponding to endosomes and larger static puncta likely corresponding to the TGN (**Movie 3**)(Liu et al., 2018). Together, these results indicate enrichment of early endocytic compartments at the apical region of neuroepithelial cells, as expected.

**Figure 1.**
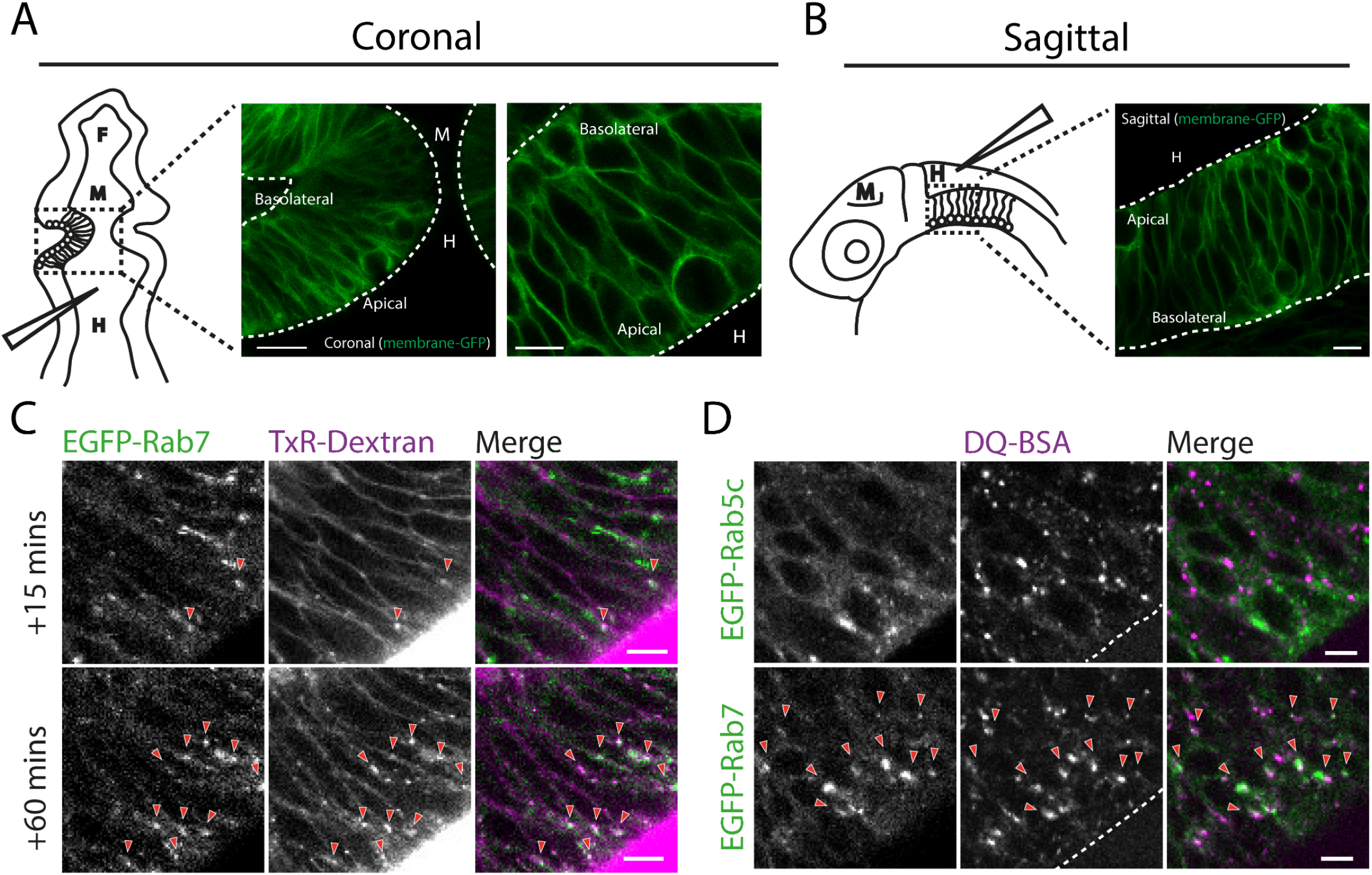
Imaging of endocytic tracer uptake and trafficking in the zebrafish neuroepithelium. Schematic of tissue organisation within the developing zebrafish brain in 1 day post-fertilisation embryos as viewed by confocal microscopy from **(A)** coronal and **(B)** sagittal perspectives. Boxed regions indicate regions imaged. Neuroepithelial cell membranes are labelled with membrane GFP. F, M and H indicate the positions of the forebrain, midbrain and hindbrain ventricle respectively with needle in black showing region injected with endocytic tracers. Dashed lines indicate apical and basolateral neuroepithelial tissue boundaries. Scale bars in A = 50 µm, 5 µm, B = 10 µm. **(C)** Hindbrain ventricle injection of TxR-Dextran accumulates in EGFP-Rab7 labelled compartments at 15 mins post-injection and to a greater degree at 60 minutes post-injection. Scale bars = 5 µm. **(D)** Hindbrain ventricle injection of DQ-BSA imaged in live embryos 1 hour post-injection shows accumulation of DQ-BSA in EGFP-Rab7 positive compartment but not EGFP-Rab5 labelled compartments. Arrowheads point to instances of co-localisation between EGFP-Rab5 or Rab7 and the indicated tracer. Scale bars = 5 µm

Injection of fluid-phase tracers into the zebrafish ventricular system has been used to gain insights into brain ventricle morphogenesis (Lowery and Sive 2005, Fame, Cortes-Campos et al. 2020), but so far not used to study endocytosis into neuroepithelial cells. We therefore first set out to assess endocytosis of fluid-phase tracers, and for this purpose injected Texas Red (TxR)-conjugated dextran into the hindbrain ventricle of zebrafish expressing the late endosome marker EGFP-Rab7 and imaged embryos live by confocal microscopy. TxR-dextran puncta were visible at 15 min post-injection, likely corresponding to its presence in endosomes, which partially overlapped with Rab7 (**Fig. 1C**). At 1 h post-injection the TxR-dextran puncta were more numerous and brighter and showed greater colocalization with Rab7, consistent with continual uptake and trafficking through late endosomes (**Fig. 1C**). We also trialled injection of the fluid phase tracer DQ-BSA, which becomes fluorescent upon proteolytic cleavage in late endosomes (Bright et al., 2016; Marwaha and Sharma, 2017), and performed live imaging. After 1 hour of uptake, DQ-BSA was present in cytoplasmic puncta, indicating effective endocytosis and delivery to late endosomes (**Movie 4**). This was confirmed by colocalization of DQ-BSA puncta with EGFP-Rab7 puncta, whereas there was little overlap with the early endosome marker EGFP-Rab5c (**Fig. 1D**). These results demonstrate that injected fluid-phase tracers undergo continual endocytosis from the ventricle and are transported to late endosomes and lysosomes. Thus, neuroepithelial cells can readily endocytose fluid-phase tracers injected into the hindbrain ventricle, which can be visualized by live imaging.

### A semi-quantitative assay for receptor-mediated endocytosis in zebrafish neuroepithelial tissue

Bulk endocytosis has been observed previously in the zebrafish neuroepithelium using the lipophilic dye FM4-64 (Clark et al., 2011), and we have also visualised fluid-phase endocytosis, as described above. However, in order to directly visualize receptor-mediated endocytosis, a specific ligand is required. We decided to use receptor-associated protein (RAP), which is a ligand for LDL receptor-related protein (LRP) family of receptors, of which LRP2 is known to be enriched on the apical pole of neuroepithelial cells (Kur et al., 2011; McCarthy et al., 2002; Spoelgen et al., 2005). Fluorescently-conjugated RAP (RAP-Cy3) was injected into the hindbrain ventricle and its endocytosis and trafficking in neuroepithelial cells assessed by live imaging. RAP endocytosis into apically-localized endosomes could be clearly visualized (**Movie 5 and 6**) with uptake detected as early as 3 minutes post-injection and the amount internalised increasing over time, as expected (**Fig. S2A**,**B** and **Movie 5**). At 3 minutes post-injection, there was significant overlap of RAP with EGFP-Rab5c, and little colocalization with EGFP-Rab7, indicating delivery to early but not late endosomes at this timepoint (**Fig. 2A-C**). At 10 and 20 minutes post-injection, the colocalization with EGFP-Rab5c remained constant, whereas it increased with EGFP-Rab7, indicating continued delivery into early endosomes and delivery of the ligand to late endosomes following its internalization and transit through early endosomes (**Fig. 2A-C**).

**Figure 2.**
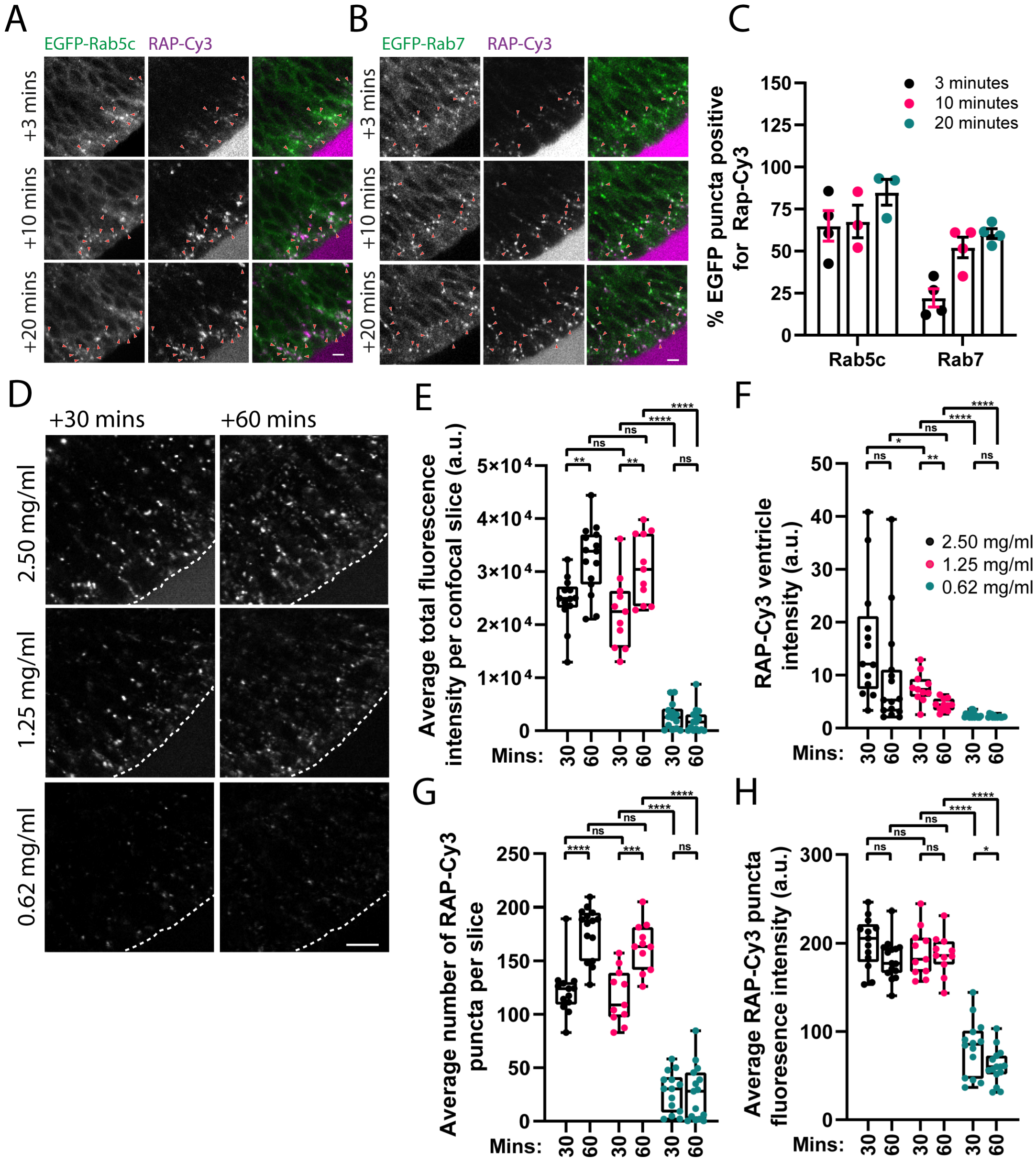
Characterisation of neuroepithelial RAP endocytosis and endocytic trafficking. Confocal microscopy images of RAP-Cy3 co-localisation with **(A)** EGFP-Rab5c and **(B)** EGFP-Rab7 in live zebrafish embryos at 3-, 10-and 20-minutes post-injection of 2.5 ng of RAP-Cy3 into the hindbrain ventricle. Arrowheads indicate co-localisation between RAP-Cy3 and Rab5c or Rab7. Scale bars = 5 µm. **(C)** Quantification of co-localisation in images between RAP-Cy3 and EGFP-Rab5c (n = 3) and EGFP-Rab7 (n = 4) at the indicated timepoints. Each datapoint represents 1 individual embryo. Error bars = S.D. **(D)** Representative live confocal microscopy images of RAP-Cy3 accumulation in neuroepithelial cells at 30 and 60 minutes post-injection of 2.5, 1.25 or 0.62 mg/ml RAP-Cy3. Scale bar = 10 µm. Quantification of **(E)** the average total fluorescence intensity per confocal slice, **(F)** RAP-Cy3 ventricle intensity at 30 or 60 minutes, **(G)** the average number of RAP-Cy3 puncta per confocal slice and **(H)** the average segmented RAP-Cy3 puncta fluorescence intensity post-hindbrain injection of the indicated concentrations of RAP-Cy3 (2.5 mg/ml, n = 15, 1.25 mg/ml, n = 11, 0.625 mg/ml, n =15). *<0.05; **<0.01; ***<0.001; ****<0.0001; ns, not significant. a.u. arbitrary units. Statistical comparisons between groups were made using students t-test.

We also observed similar co-localisation kinetics with RAP and CLC to those for Rab5c, consistent with the presence of RAP in early endocytic intermediates (**Fig. S3A**,**C**). Lower levels of co-localisation between RAP and EGFP-Rab11a at 10 and 20 minutes post-injection could also be observed, suggesting some delivery of RAP to recycling endosomes at these timepoints (**Fig. S3B**,**C**). At 1 h post-injection, RAP was present in larger puncta that exhibited dynamic bidirectional movement through the neuroepithelial cell cytoplasm and occasionally in elongated tubular structures oriented along the apicobasal axis (**Movie 7** and **Fig. S2C**). There was a high degree of overlap with LAMP1-GFP, indicating these structures correspond to lysosomes (**Movie 8**). Interestingly, the lysosomes appeared to undergo fusion and tubulation/fission events, and RAP could often be seen in these intermediates (**Movie 8**). Thus, as with the fluid-phase markers DQ-BSA and TxR dextran, the receptor ligand RAP is also endocytosed by neuroepithelial cells and follows a similar itinerary, accumulating firstly in early endosomes before being transported to late endosomal and lysosomal compartments.

To quantify RAP endocytosis, we titrated the concentration injected and measured a number of parameters based on the amount of RAP fluorescence within the neuroepithelial tissue. Injection of 2.5 mg/ml and 1.25 mg/ml concentrations of RAP into the hindbrain ventricle yielded higher total fluorescence intensity values per area of tissue quantified at 30-and 60-minutes post injection, compared to injection of 0.625 mg/ml RAP (**Fig. 2D,E**). Due to the presence of excess RAP in the ventricle for prolonged periods of time post-injection (**Fig. 2F**), injection of 2.5 mg/ml or 1.25 mg/ml RAP led to an increase in the amount of tissue fluorescence from 30 to 60 minutes (**Fig. 2D,E**). This increase was not seen using 0.625 mg/ml RAP (**Fig. 2D,E**) likely due to depletion of the limited amount of ligand in the ventricle **(Fig. 2F)**. Similarly, as a likely result of RAP accumulating in endosomal and lysosomal compartments over time, increasing concentrations of RAP lead to an increase in the number of detectable puncta within neuroepithelial cells (**Fig. 2G**). As an additional quantitative parameter of RAP endocytosis, we divided the total fluorescence intensity at each concentration by the total number of segmented puncta to give an average puncta intensity value. As expected, the average puncta intensity was lower for 0.625 mg/ml RAP compared to the higher concentrations (**Fig. 2H**). Collectively, these results demonstrate that RAP endocytosis at the neuroepithelium can be quantified and reliably detect the anticipated differences in endocytic uptake.

### *ocrl* mutant embryos display disrupted neuroepithelial RAP endocytosis and trafficking

We next applied the RAP uptake assay to an *ocrl* mutant zebrafish line to examine whether neuroepithelial RAP endocytosis and trafficking is impaired in the absence of Ocrl. Ocrl is a inositol-5-phosphatase that acts on pools of PI(4,5)P_2_ coupled to membrane remodelling events at multiple stages of the endocytic pathway, which includes clathrin-mediated endocytosis (Choudhury et al., 2009; Erdmann et al., 2007; Nandez et al., 2014), receptor recycling from endosomes (Choudhury et al., 2005; van Rahden et al., 2012; Vicinanza et al., 2011) and lysosome fusion (De Leo et al., 2016; Wang et al., 2021; Zhang et al., 1998). Mutations in *OCRL* are responsible for Lowe syndrome and Dent 2 disease, X-linked multi-systemic disorders that predominantly affect the eyes, kidney and central nervous system (Bokenkamp and Ludwig, 2016). *ocrl* mutant zebrafish have previously been shown to display multiple CNS phenotypes matching those seen in Lowe syndrome, including cystic brain lesions within white matter regions and increased sensitivity to febrile seizures during early development (Ramirez et al., 2012). These phenotypes were potentially linked to altered rates of proliferation and apoptosis within the zebrafish CNS at 1-day post-fertilisation (dpf) (Ramirez et al., 2012). As neuroepithelial cells are the major proliferative cell type found in the brain during the early stages of neurogenesis, we hypothesised that loss of Ocrl function may perturb endocytic trafficking within neuroepithelial cells and contribute to the aberrant CNS development seen in o*crl* mutant zebrafish.

For RAP uptake experiments we used an *ocrl* mutant (sa11582) obtained from the zebrafish mutation project (Kettleborough et al., 2013) containing a G to T mutation in a splice acceptor site at the boundary of intron 16-17 and exon 17, which encodes part of the *ocrl* ASH domain (**Fig. S4A**). This mutation abolishes a SpeI restriction enzyme site present in the wild-type zebrafish *ocrl* genomic sequence, which we could use for genotyping purposes (**Fig. S4B**). Embryos carrying this mutation showed a complete loss of Ocrl protein, potentially due to nonsense mediated decay of a faulty *ocrl* transcript, and also displayed a noticeable reduction in brain size at the same developmental stage as wild-type controls, consistent with phenotypes seen previously in a different *ocrl* mutant zebrafish line (Ramirez et al., 2012) (**Fig. S4C**,**D**).

Endocytosis of RAP was assessed in *ocrl* mutant embryos at 30 minutes following injection of 2.5 mg/ml RAP, as at this timepoint and concentration, RAP has not been depleted from the ventricle and both early and late endosomes are labelled. Live imaging showed that RAP uptake was reduced in *ocrl* mutants, with lower total fluorescence and number of detectable puncta in the neuroepithelial cells (**Fig. 3A-C**). Quantitation revealed a slight reduction in the average intensity of RAP puncta in *ocrl* mutant embryos, but this was not significant (**Fig. 3D**), and there was also a slight but not significant increase in the amount of RAP remaining in the ventricle at 30 minutes (**Fig. 3E**). RAP also did not appear to traffic as deep into the neuroepithelial cells in the *ocrl* mutant, suggesting delayed transit along the endocytic pathway (**Fig. 3A**). In support of an effect on endosomal trafficking, EEA1-positive early endosomes were enlarged in *ocrl* mutant embryos, and similarly to RAP, these appeared to more apically localized compared to WT embryos (**Fig. 3F**,**G**). Additionally, total Lrp2 abundance was reduced in the neuroepithelial cells of the *ocrl* mutant, with a striking reduction at the apical surface, consistent with a defect in its endosomal trafficking, most likely in recycling back to the apical pole (**Fig. 3H**,**I**). A similar reduction in Lrp2 abundance was reported in *ocrl*-deficient renal proximal tubule cells and has been attributed to defects in receptor recycling (Festa et al., 2019; Oltrabella et al., 2015).

**Figure 3.**
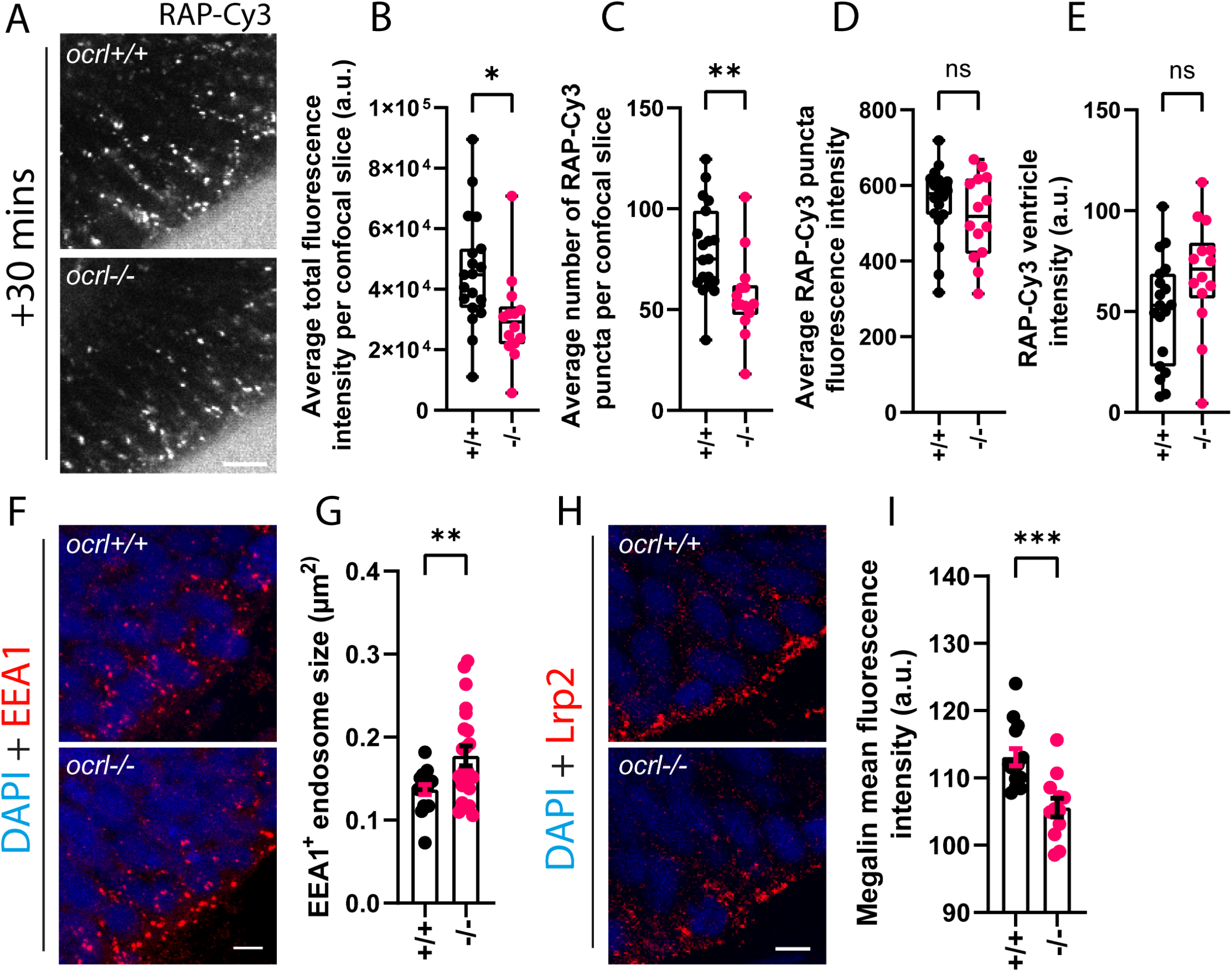
Ocrl regulates receptor recycling and trafficking of endocytic ligands in zebrafish neuroepithelial cells. **(A)** Representative confocal images of endocytic uptake of RAP-Cy3 30 minutes post-hindbrain ventricle injection in live *ocrl* mutant (n = 14) embryos or WT controls (n = 19). Scale bar = 10 µm. Quantification of **(B)** the average total fluorescence intensity per confocal slice, **(C)** average number of RAP-Cy3 puncta per confocal slice, **(D)** average RAP-Cy3 puncta fluorescence intensity, and **(E)** RAP-Cy3 ventricle intensity at 30 minutes post-hindbrain injection of RAP-Cy3 in WT and *ocrl* mutant embryos. **(F)** Representative confocal microscopy images of fixed tissue sections from WT (n = 17) or *ocrl* mutant (n = 22) embryos stained with antibodies against EEA1 and **(G)** quantification of EEA1 positive endosome size in WT and *ocrl* mutant embryos. Scale bar = 5 µm. **(H)** Representative confocal microscopy images of fixed tissue sections through the zebrafish hindbrain from WT (n = 14) or *ocrl* mutant (n = 12) embryos stained with antibodies against Lrp2. **(I)** Quantification of the mean Lrp2 fluorescence intensity from WT or *ocrl* mutant embryos. Scale bar = 5 µm. *<0.05; **<0.01; ***<0.001; ****<0.0001; ns, not significant. a.u. arbitrary units. Statistical comparisons between groups were made using students t-test.

### Ocrl localises to endocytic compartments in the zebrafish neuroepithelium

We next wanted to localize Ocrl in the zebrafish neuroepithelium. Live imaging of embryos transiently expressing an EGFP tagged version of the brain specific Ocrl isoform, Ocrla (Johnson et al., 2003), revealed Ocrl-containing puncta that often co-localised with endocytosed RAP and to a much lesser extent DQ-BSA, although high cytosolic levels of EGFP-Ocrla made visualisation of the puncta difficult (**Fig. 4A**). To further investigate whether Ocrl localises to endocytic compartments in zebrafish neuroepithelial cells, we co-labelled tissue sections through the neuroepithelium of 1 dpf embryos with various endocytic and organelle markers (**Fig. 4B,C**). Fixation and permeabilization of tissue sections removed a significant fraction of the cytosolic EGFP signal revealing multiple punctate structures throughout the cytoplasm. The vast majority of these co-localised with the Golgi marker Golgin-84, consistent with the reported localisation of Ocrl to the Golgi apparatus (Hyvola et al., 2006). A significant amount of overlap was also seen between EGFP-Ocrla and both EEA1 and Lrp2, indicating that Ocrl is recruited to early endosomal compartments within neuroepithelial cells that also likely contain Lrp2 (**Fig. 4B,C**). A low amount of co-localisation of EGFP-Ocrla with the late endosomal and lysosome marker LAMP1 was also observed. Together, these results provide evidence that Ocrl functions within the endosomal system of neuroepithelial cells to maintain normal receptor traffic.

**Figure 4.**
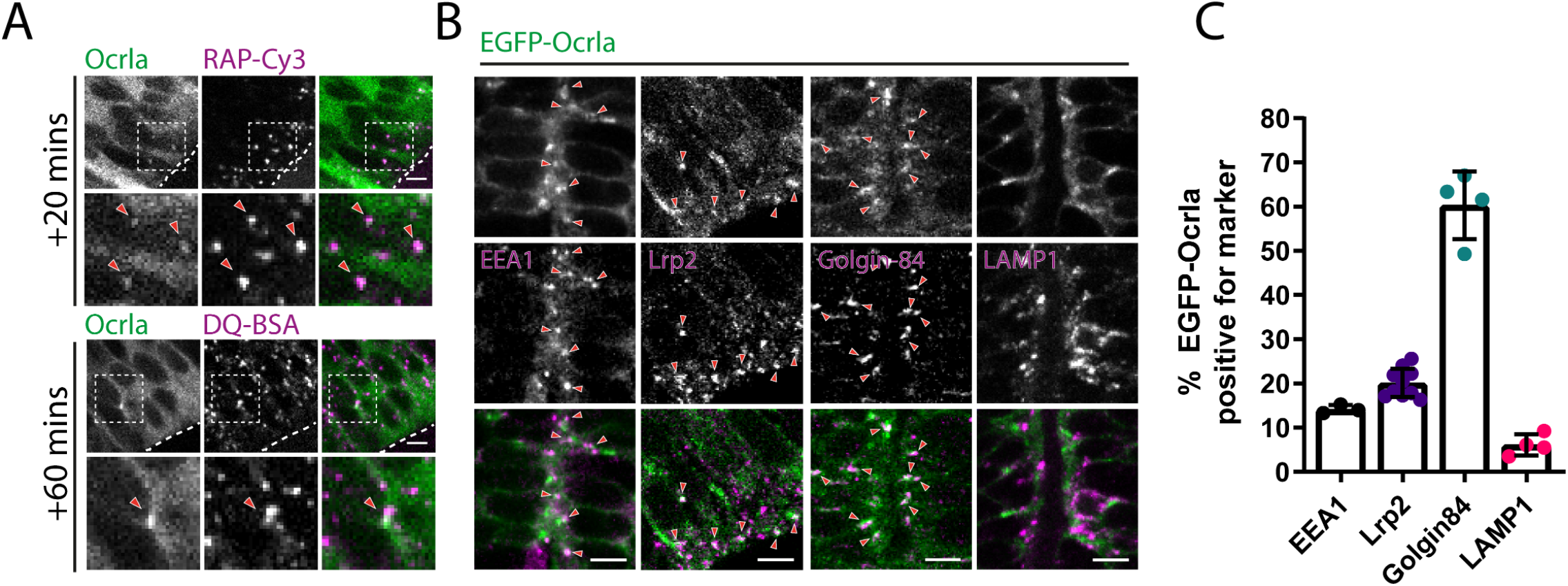
Ocrl localises to early endocytic compartments in neuroepithelial cells. **(A)** Representative confocal images of live 28 hpf zebrafish embryos expressing EGFP-Ocrla after hindbrain ventricle injection of RAP-Cy3 or DQ-BSA. Arrowheads point to co-localisation between Ocrla signal and RAP-Cy3 or DQ-BSA. Scale bar = 5 µm. **(B)** Representative images of fixed tissue sections through the hindbrain of embryos expressing EGFP-Ocrla, stained with antibodies against either EEA1, Lrp2, Golgin-84 or LAMP1. Arrowheads indicate instances of co-localisation between Ocrla and the indicated marker. Scale bar = 5 µm. **(C)** Quantification of co-localisation between Ocrla puncta and EEA1 (n = 3), Lrp2 (n = 10), Golgin-84 (n = 4) or LAMP1 (n = 4) on fixed tissue sections. Each data point represents one individual embryo. Error bars = S.D.

### Neuroepithelial RAP endocytosis can occur independently of Lrp2

In the zebrafish pronephros, endocytosis of RAP is dependent upon Lrp2, which is highly abundant at the apical membrane of the proximal tubule epithelium. In this tissue, RAP can therefore be used as a readout for Lrp2 endocytosis. However, RAP can bind to additional members of the LRP family of endocytic receptors other than LRP2 (Andersen et al., 2003; Battey et al., 1994; Kounnas et al., 1992). We therefore wanted to know whether RAP uptake in the zebrafish neuroepithelium is dependent on Lrp2 or other LRP family members and towards this end first decided to localize Lrp2 in the zebrafish neuroepithelium. For this purpose, co-localization of Lrp2 was performed on cryosections of neuroepithelial tissue prepared from embryos expressing EGFP tagged Rab5c, Rab11a or LAMP1. In agreement with previous studies performed in mice and zebrafish (Christ et al., 2012; Kur et al., 2011), Lrp2 was abundant at the apical pole of neuroepithelial cells, localizing to puncta at the apical membrane and proximal to it (**Fig. 5A**). The Lrp2 containing puncta strongly co-localised with Rab11a (80%) and to a lesser extent with Rab5c (40%) (**Fig. 5A,B**), consistent with the transit of Lrp2 through early endosomes and its recycling via Rab11 recycling endosomes (Nagai et al., 2003; Perez Bay et al., 2016; Shah et al., 2013). A comparatively small amount of Lrp2 also co-localised with late endosomal and lysosomal compartments labelled with LAMP1-GFP, again in agreement with previous results (Christ et al., 2012).

**Figure 5.**
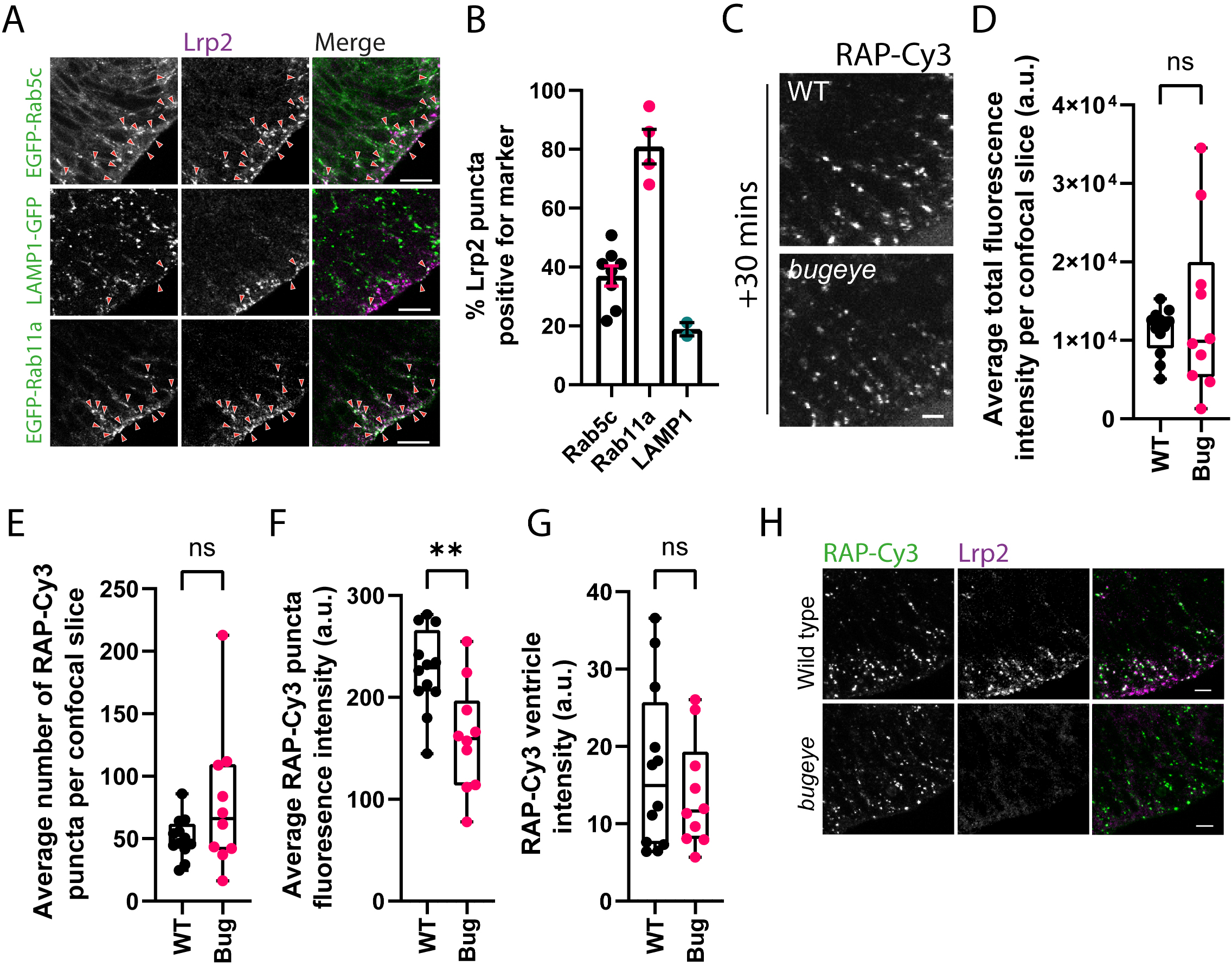
Analysis of neuroepithelial endocytic uptake of RAP in *lrp2* knockout embryos. **(A)** Tissue sections collected from embryos transiently expressing EGFP-tagged Rab5c, Rab11a of LAMP1 at 28 hpf and stained with antibodies against Lrp2 to visualise Lrp2 localisation with the zebrafish neuroepithelium. Arrowheads point to examples of co-localisation between Lrp2 and the indicated marker. Scale bars = 10 µm **(B)** Quantification of Lrp2 co-localisation with EGFP tagged Rab5c (n = 8), Rab11a (n = 4) or LAMP1 (n = 2). Each datapoint represents quantification from one individual embryo. EGFP tagged constructs were expressed transiently. Error bars = S.D. **(C)** Assessment of endocytic uptake of RAP in *lrp2* knockout embryos. Representative confocal microscopy images of live WT or *lrp2* null zebrafish embryos following hindbrain ventricle injection of RAP-Cy3 imaged 30 minutes post-injection. Scale bar = 5µm. Quantification of **(D)** the average total fluorescence intensity per confocal slice, **(E)** the average number of RAP-Cy3 puncta per confocal slice, **(F)** the average RAP-Cy3 puncta fluorescence intensity, and **(G)** RAP-Cy3 ventricle intensity at 30 post-hindbrain injection of RAP-Cy3 in WT (n = 12) and *lrp2* mutant (n = 10) embryos. **(H)** Tissue sections through the zebrafish hindbrain from WT or *lrp2* mutant embryos injected with RAP-Cy3, fixed 30 minutes post-injection and stained with antibodies against Lrp2. Scale bar = 10 µm. *<0.05; **<0.01; ***<0.001; ****<0.0001; ns, not significant. a.u. arbitrary units. Statistical comparisons between groups were made using students t-test.

To determine the dependence of RAP endocytosis on Lrp2 in the zebrafish neuroepithelium, *bugeye lrp2* knockout embryos (Veth et al., 2011) were injected into the hindbrain ventricle with RAP, and uptake was quantified in live embryos at 30 minutes post-injection. Surprisingly, neuroepithelial RAP uptake was efficient in the *lrp2* knockout embryos (**Fig. 5C**). There was no significant difference in either the total fluorescence intensity, number of RAP puncta or ventricle intensity in between WT and *lrp2* null embryos (**Fig. 5D**,**E**,**G**). However, the average RAP puncta intensity in *lrp2* null embryos was marginally lower than WT controls (**Fig. 5F**), consistent with a slight reduction in RAP endocytosis. RAP uptake was also visualised by staining of WT and *bugeye* tissue sections taken from embryos injected with RAP and fixed at 30 minutes post-injection. This confirmed that *bugeye* embryos, despite completely lacking Lrp2 at the apical surface, endocytosed RAP into the neuroepithelial tissue at similar levels to WT controls (**Fig. 5H**). Endocytosis of RAP into the zebrafish neuroepithelium would therefore appear to depend on additional receptors present at the apical surface of neuroepithelial cells.

## Discussion

In this manuscript, we describe methods to assess both fluid-phase and receptor-mediated endocytosis in the developing zebrafish brain. These methods involve injection of tracer into the brain ventricle and live or fixed imaging to assess endocytosis into the surrounding neuroepithelium. Our work builds upon previous studies in zebrafish using FM4-64 to visualise bulk endocytosis (Clark et al., 2011), and more recently, an antibody uptake approach to visualise endocytosis of the membrane-bound Notch ligand DeltaD (Zhao et al., 2021). The latter represents an assay for receptor-mediated endocytosis, albeit of an atypical membrane-associated receptor ligand. Antibody uptake assays require the availability of an antibody to the ligand or the extracellular region of the receptor of interest, and antibody binding may alter the trafficking kinetics of the proteins under investigation. In our experiments, RAP was used a more conventional soluble ligand for receptor-mediated uptake. It binds to LRP family receptors (Kounnas et al., 1992), discussed further below, and can be easily followed by microscopy to reveal its sorting and delivery to lysosomes. Importantly, its uptake and endocytic trafficking can be quantified. Our results therefore show that quantitative assessment of endocytosis in the neuroepithelium is feasible, which provides the basis for a standardised protocol to measure this process in vivo under different experimental conditions.

The ability to perform live imaging of the zebrafish embryo neuroepithelium allows not only assessment of ligand uptake, but also the dynamics of trafficking within the endosomal system. Moreover, the ability to express transgenes in zebrafish embryos allows for the dual imaging of endocytic cargoes, receptors, compartment markers or trafficking machinery within the neuroepithelium (Beis and Stainier, 2006; Vacaru et al., 2014). We could exploit this property to co-visualise RAP with markers of different endosome types and lysosomes to assess traffic of the ligand through the endocytic pathway. Hence, the zebrafish embryo neuroepithelium is a highly tractable system for studying endocytic traffic in vivo. One limitation is that injection of tracers into the ventricle only allows for assessment of endocytosis at the apical pole, but, as for other epithelial cells, apical endocytosis appears to be a highly active process in neuroepithelial cells (Aaku-Saraste et al., 1996; Aaku-Saraste et al., 1997; Zhao et al., 2021), making it suitable for the imaging approaches described.

We applied the RAP uptake assay to study the role of Ocrl in endocytic traffic in the developing zebrafish neuroepithelium and observed several endocytic defects in *ocrl* null embryos. There was reduced RAP uptake into the neuroepithelial cells, consistent with a role for Ocrl in clathrin-mediated endocytosis at the plasma membrane, as shown in previous in vitro studies (Choudhury et al., 2005; Choudhury et al., 2009; Erdmann et al., 2007; Nandez et al., 2014). It would also align with our previous work showing that clathrin-binding is important for Ocrl function in the developing brain (Ramirez et al., 2012). Our imaging experiments also indicated reduced movement of RAP from the sub-apical region to deeper within cells, suggesting reduced progression along the endocytic pathway, and we also observed that early endosomes were enlarged. These effects can be explained by defective trafficking at early endosomes, as reported previously for loss of Ocrl and its interaction partners in vitro (Choudhury et al., 2005; Erdmann et al., 2007; Festa et al., 2019; Noakes et al., 2011; van Rahden et al., 2012; Vicinanza et al., 2011). The reduced abundance of Lrp2 at the apical pole in *ocrl* mutants would be consistent with reduced recycling from endosomes, with a similar phenomenon reported previously in renal proximal tubule cells (Festa et al., 2019; Oltrabella et al., 2021; Oltrabella et al., 2015) Thus, it is likely that Ocrl operates at multiple stages of the endocytic pathway in the neuroepithelium. Further studies will be necessary to determine the precise steps and cargos regulated by Ocrl activity within the neuroepithelial endocytic pathway.

Surprisingly, RAP endocytosis at the neuroepithelium was only marginally affected by loss of Lrp2. In zebrafish, the *lrp2* gene has been duplicated, to generate *lrp2a* and *lrp2b* forms, but only *lrp2a* is lost in the *bugeye* mutant (Kur et al., 2011). This raises the possibility of Lrp2b being responsible for RAP uptake in *bugeye* embryos. However, previous work has shown that *lrp2b* is expressed at much lower levels than *lrp2a*, and that it does not functionally compensate for the loss of *lrp2a* during development (Kur et al., 2011). This suggests that another RAP-binding receptor is present at the apical surface of the neuroepithelium. The identity of this receptor is currently unclear. RAP can bind to other members of the LRP receptor family and several of these (LRP1, ApoER2, VLDLR) are expressed in radial glia cells (Auderset et al., 2016a; Bres et al., 2020; Luque et al., 2003), a related neuroepithelial cell type that also have an apical membrane exposed to the ventricle (Gotz and Huttner, 2005; Taverna et al., 2014). However, these receptors appear to be localised to the basolateral membrane of the radial glia cells (Auderset et al., 2016a; Auderset et al., 2016b; Boggild et al., 2018; Luque et al., 2003), and are unlikely to be the receptor for injected RAP. LRP5 and 6, which play an important role in Wnt signalling during many developmental processes, including neural development (Angers and Moon, 2009; Gray et al., 2013; Jeong et al., 2014), can also bind RAP, albeit with lower affinity than other members of the LRP family (Li et al., 2005). As LRP5 and LRP6 have been reported to be present at the apical surface of other epithelial cell types, they are also potentially present at the apical membrane of neuronal precursors and hence may play a role in RAP uptake at the neuroepithelium (Takita and Seko, 2020; Yamamoto et al., 2017). Further studies are necessary to identify the RAP receptor at the apical membrane of the zebrafish neuroepithelium.

It is interesting to note that loss of Lrp2 does not affect neural development in zebrafish, whereas in mammals it causes forebrain malformation, due to defects in SHH endocytosis and morphogen gradient formation (Christ et al., 2012; Kur et al., 2011). This difference may reflect different requirements for SHH signalling between mammals and teleosts, or perhaps differential requirements for LRP family receptors in endocytosis and developmental signalling between the two classes of vertebrates (Willnow and Christ, 2017). Uncovering the complement of receptors that mediate zebrafish neuroepithelial RAP endocytosis may therefore provide further insights into whether and how LRP family members contribute to neural development in zebrafish and possibly in mammals too (Christ et al., 2012; Kur et al., 2011). Using more specific ligands for the different LRP family members and specific knockouts of these family members should prove informative in this regard.

## Materials and methods

### Antibodies

Antibodies used as part of this study were sheep anti-zebrafish OCRL (WB 1:200) (Ramirez et al., 2012), sheep anti-Golgin-84 (IF 1:1000) (Diao et al., 2003), goat anti-EEA1 (IF 1:400) (SantaCruz Biotech), rabbit anti-LAMP1 (IF 1:200) (Abcam24170) and rabbit anti-megalin/Lrp2 (IF 1:100). The anti-megalin/Lrp2 antibody was a kind gift from Dr Michle Marino, University of Pisa, Italy.

### Zebrafish husbandry and strains

Zebrafish were raised and maintained by the University of Manchester Biological Services Facility in accordance with UK Animals Act 1986. Embryos used as WT controls were from the AB Notts strain unless otherwise stated. Transgenic GFP-Rab5c, GFP-Rab7, GFP-Rab11a and *lrp2* null *bugeye* mutant zebrafish were kind gifts courtesy of Professor Brian Link, Medical College of Wisconsin and described previously (Clark et al., 2011; Veth et al., 2011). The *ocrl* mutant sa11582 was generated by ENU mutagenesis as part of the zebrafish mutation project (Kettleborough et al., 2013). *ocrl* mutant sa11582 was backcrossed to the wild type AB Notts line to generate *ocrl*^+/-^ heterozygotes before in crossing to obtain a pool of fish homozygous for the sa11582 *ocrl* allele. From the same in crosses, a pool of *ocrl*^+/+^ fish was also isolated and maintained for use as sibling controls for all experiments investigating Ocrl function.

### Molecular biology and mRNA synthesis

GFP tagged rat LAMP1 and zebrafish Rab5c, Rab7 and Rab11a were cloned into pcGlobin (Ro et al., 2004) between EcoRI and EcoRV sites using standard molecular biology techniques. Primers for these constructs are available upon request. mRNA for transient expression of EGFP tagged fusion proteins were made by linearisation of plasmid DNA followed by mRNA synthesis and purification using either T7 or SP6 mMessage transcription kit (Ambion). For transient expression, one cell stage embryos were injected with 1nl of the purified mRNA solutions at the indicated concentrations: EGFP-Ocrla 750 ng/µl, GFP-CLC 250 ng/µl, GFP-Rab5c 250 ng/µl, GFP-Rab7 250 ng/µl, GFP-Rab11a 250 ng/µl, GFP-LAMP1 250ng/µl.

### Genotyping of *ocrl* mutant adult zebrafish

For genotyping of *ocrl* mutant adult zebrafish a PCR fragment spanning the SpeI restriction site in the *ocrl* gene was amplified from genomic DNA prepared from adult fish as previously described (Wilkinson et al., 2013). The resulting 720 bp fragment was digested using SpeI restriction enzyme (New England Biolabs) and the digested product resolved by gel electrophoresis. Fish were classified as *ocrl*^+/+^, or *ocrl*^−/-^ by the presence of a double band indicating digestion (*ocrl*^+/+^) or a single band indicating resistance to digestion (*ocrl*^−/-^).

### Western blotting and antibodies

Embryos at the appropriate stage were culled by incubation with 0.2 mg/ml MS222 (Sigma) for >2h. To remove yolk proteins, 500μl Ginsburg fish Ringer’s solution (110 mM NaCl, 3.5 mM KCl, 2.7 mM CaCl_2_._2_H_2_O, 2.3 mM NaHCO_3_, 10 mM Tris pH 8.5) was added to embryos and the yolk sac disrupted mechanically through pipetting (Link et al., 2006). Following two washes in Ginsburg, embryos were pelleted at 300 x g for 1 minute and Ringer’s solution removed. An appropriate volume of RIPA buffer (20 mM Tis-HCl, 150 mM NaCl, 1 mM EGTA, 1% (v/v) NP-40 substitute, 1% (w/v) sodium deoxycholate) was then added and embryos homogenised in a 1.5 ml microcentrifuge tube using a pestle. Lysates were centrifuged for five minutes at 15,000 x g to pellet debris and transferred to a fresh 1.5 ml tube. 6 μl of lysate was removed and protein concentration determined by BCA assay. Lysates were resolved by SDS-PAGE, transferred to nitrocellulose membranes, blocked in 5% (w/v) milk dissolved in PBS & 0.1% Tween (PBSTw) before being probed overnight with primary antibody. Secondary antibody incubations were performed the next day for one hour following three washes in PBSTw to remove unbound primary antibody. Following secondary antibody incubation, membranes were washed a further three times in PBST for 5 minutes each before imaging by chemiluminescence on a BioRad gel doc imaging station.

### Cryosections and immunofluorescence staining of tissue sections

Embryos were fixed in 4% (v/v) formaldehyde in PBS (pH 7.4) at 4°C overnight or at room temperature for 2-3 hours before being rinsed three times in PBS & 0.1% (v/v) Triton X-100 (PBST) and dehydrated for a minimum of 30 minutes in methanol at -20°C. Embryos were then rehydrated in descending 75%, 50% and 25% concentrations of methanol with PBST, rinsed three times in PBST and embedded overnight in gelatin (15% (v/v) fish skin gelatin, 30% (w/v) sucrose in PBS). 12 μm sections were then cut using a Leica CM3050 cryostat tissue sections collected on coated Superfrost plus slides (Thermofisher). Sectioned material was dried for 1 hour at room temperature before being submerged in 100% acetone at room temperature for 2 minutes and rehydrated in PBS for 10 minutes. To reduce non-specific binding of antibodies, slides were incubated with blocking solution (PBST & 10% (v/v) donkey serum) at room temperature in a humidified atmosphere. Following blocking, primary antibodies in blocking solution were added to slides and incubated for 2 hours at room temperature. After primary antibody incubation, slides were rinsed in PBST 6 times for five minutes each. Secondary antibodies in blocking solution were then added to slides for 2 hours at room temperature. Slides were again washed 6 times in PBST for 5 mins each before mounting of cover slips using Mowiol 4-88.

### Injection and live imaging of the zebrafish neuroepithelium

To facilitate live imaging, N-phenylthiourea (PTU) was added to the water at 24 hours post-fertilisation to inhibit melanin synthesis. For injection of endocytic tracers into the hindbrain ventricle, embryos were first anaesthetised in 0.2 mg/ml MS222 before being oriented in an agarose dish for injection and imaging as previously described (Chang and Sive, 2012). Approximately 1 nl of each tracer was injected into the hindbrain ventricle close to the midbrain hindbrain boundary before embryos were imaged live on either a Leica SP5 or SP8 confocal microscope with a 63x water immersion objective. Per injection, a volume of 50 µm per embryo was imaged at 1.5 µm intervals beginning at the roofplate of the dorsal MHB.

### Quantification of RAP Cy3 endocytosis

Images were quantified in FIJI (Schindelin et al., 2012). In RAP injected embryos, an equivalent 40 µm^2^ area of tissue within one lobe of the midbrain hindbrain boundary was first selected. A standardised threshold was then applied to all images to exclude background fluorescence from the quantification. RAP-Cy3 puncta were segmented within each confocal plane using the analyse particles function in Image J. For each confocal plane captured and analysed, the average total fluorescence intensity and puncta number was calculated. The average total fluorescence intensity and number of puncta for each embryo injected was then calculated by taking an average total fluorescence intensity and puncta number from each confocal plane over a minimum range of 10 confocal slices starting from where the first complete cell bodies of MHB neuroepithelial cells are visible. To prevent incorporating fluorescence remaining in the ventricle as part of the quantification, segmented puncta above a size of 5 µm were excluded from measurements used to calculate averages for each parameter. Ventricle intensity measurements were made in the first confocal plane from the dorsal MHB where RAP fluorescence could be visualised in the ventricle.

## Statistical analysis

All statistical comparisons between groups were made in GraphPad Prism 8 or 9 using an unpaired students *t*-test. Bars and error bars in data presented as a bar chart represent the mean value ± standard deviation. Values from individual embryos are overlaid. For box and whisker plots, boxes represent upper and lower quartile values with median values indicated by a line inside the box. Whiskers demonstrate the highest and lowest value within each treatment group or genotype. Data presented are from a minimum of 2 independent experiments. N numbers represent the total number of embryos analysed from all independent experiments.

## Acknowledgements

We are grateful to the University of Manchester bioimaging facility for help with confocal imaging experiments.

## References

Aaku-Saraste, E., Hellwig, A. and Huttner, W. B. (1996). Loss of occludin and functional tight junctions, but not ZO-1, during neural tube closure--remodeling of the neuroepithelium prior to neurogenesis. Dev Biol 180, 664–79.

Aaku-Saraste, E., Oback, B., Hellwig, A. and Huttner, W. B. (1997). Neuroepithelial cells downregulate their plasma membrane polarity prior to neural tube closure and neurogenesis. Mech Dev 69, 71–81.

Andersen, O. M., Benhayon, D., Curran, T. and Willnow, T. E. (2003). Differential binding of ligands to the apolipoprotein E receptor 2. Biochemistry 42, 9355–64.

Angers, S. and Moon, R. T. (2009). Proximal events in Wnt signal transduction. Nat Rev Mol Cell Biol 10, 468–77.

Auderset, L., Cullen, C. L. and Young, K. M. (2016a). Low Density Lipoprotein-Receptor Related Protein 1 Is Differentially Expressed by Neuronal and Glial Populations in the Developing and Mature Mouse Central Nervous System. PLoS One 11, e0155878.

Auderset, L., Landowski, L. M., Foa, L. and Young, K. M. (2016b). Low Density Lipoprotein Receptor Related Proteins as Regulators of Neural Stem and Progenitor Cell Function. Stem Cells Int 2016, 2108495.

Battey, F. D., Gafvels, M. E., FitzGerald, D. J., Argraves, W. S., Chappell, D. A., Strauss, J. F., 3rd and Strickland, D. K. (1994). The 39-kDa receptor-associated protein regulates ligand binding by the very low density lipoprotein receptor. J Biol Chem 269, 23268–73.

Beis, D. and Stainier, D. Y. (2006). In vivo cell biology: following the zebrafish trend. Trends Cell Biol 16, 105–12.

Boggild, S., Molgaard, S., Glerup, S. and Nyengaard, J. R. (2018). Highly segregated localization of the functionally related vps10p receptors sortilin and SorCS2 during neurodevelopment. J Comp Neurol 526, 1267–1286.

Bokenkamp, A. and Ludwig, M. (2016). The oculocerebrorenal syndrome of Lowe: an update. Pediatr Nephrol 31, 2201–2212.

Bres, E. E., Safina, D., Muller, J., Bedner, P., Yang, H., Helluy, X., Shchyglo, O., Jansen, S., Mark, M. D., Esser, A. et al. (2020). Lipoprotein receptor loss in forebrain radial glia results in neurological deficits and severe seizures. Glia 68, 2517–2549.

Bright, N. A., Davis, L. J. and Luzio, J. P. (2016). Endolysosomes Are the Principal Intracellular Sites of Acid Hydrolase Activity. Curr Biol 26, 2233–45.

Chae, T. H., Kim, S., Marz, K. E., Hanson, P. I. and Walsh, C. A. (2004). The hyh mutation uncovers roles for alpha Snap in apical protein localization and control of neural cell fate. Nat Genet 36, 264–70.

Chang, J. T. and Sive, H. (2012). An assay for permeability of the zebrafish embryonic neuroepithelium. J Vis Exp, e4242.

Choudhury, R., Diao, A., Zhang, F., Eisenberg, E., Saint-Pol, A., Williams, C., Konstantakopoulos, A., Lucocq, J., Johannes, L., Rabouille, C. et al. (2005). Lowe syndrome protein OCRL1 interacts with clathrin and regulates protein trafficking between endosomes and the trans-Golgi network. Mol Biol Cell 16, 3467–79.

Choudhury, R., Noakes, C. J., McKenzie, E., Kox, C. and Lowe, M. (2009). Differential clathrin binding and subcellular localization of OCRL1 splice isoforms. J Biol Chem 284, 9965–73.

Christ, A., Christa, A., Kur, E., Lioubinski, O., Bachmann, S., Willnow, T. E. and Hammes, A. (2012). LRP2 is an auxiliary SHH receptor required to condition the forebrain ventral midline for inductive signals. Dev Cell 22, 268–78.

Clark, B. S., Miesfeld, J. B., Flinn, M. A., Collery, R. F. and Link, B. A. (2020). Dynamic Polarization of Rab11a Modulates Crb2a Localization and Impacts Signaling to Regulate Retinal Neurogenesis. Front Cell Dev Biol 8, 608112.

Clark, B. S., Winter, M., Cohen, A. R. and Link, B. A. (2011). Generation of Rab-based transgenic lines for in vivo studies of endosome biology in zebrafish. Dev Dyn 240, 2452–65.

Coumailleau, F., Furthauer, M., Knoblich, J. A. and Gonzalez-Gaitan, M. (2009). Directional Delta and Notch trafficking in Sara endosomes during asymmetric cell division. Nature 458, 1051–5.

De Leo, M. G., Staiano, L., Vicinanza, M., Luciani, A., Carissimo, A., Mutarelli, M., Di Campli, A., Polishchuk, E., Di Tullio, G., Morra, V. et al. (2016). Autophagosome-lysosome fusion triggers a lysosomal response mediated by TLR9 and controlled by OCRL. Nat Cell Biol 18, 839–850.

Diao, A., Rahman, D., Pappin, D. J., Lucocq, J. and Lowe, M. (2003). The coiled-coil membrane protein golgin-84 is a novel rab effector required for Golgi ribbon formation. J Cell Biol 160, 201–12.

Emery, G. and Knoblich, J. A. (2006). Endosome dynamics during development. Curr Opin Cell Biol 18, 407–15.

Erdmann, K. S., Mao, Y., McCrea, H. J., Zoncu, R., Lee, S., Paradise, S., Modregger, J., Biemesderfer, D., Toomre, D. and De Camilli, P. (2007). A role of the Lowe syndrome protein OCRL in early steps of the endocytic pathway. Dev Cell 13, 377–90.

Fame, R. M., Cortes-Campos, C. and Sive, H. L. (2020). Brain Ventricular System and Cerebrospinal Fluid Development and Function: Light at the End of the Tube: A Primer with Latest Insights. Bioessays 42, e1900186.

Festa, B. P., Berquez, M., Gassama, A., Amrein, I., Ismail, H. M., Samardzija, M., Staiano, L., Luciani, A., Grimm, C., Nussbaum, R. L. et al. (2019). OCRL deficiency impairs endolysosomal function in a humanized mouse model for Lowe syndrome and Dent disease. Hum Mol Genet 28, 1931–1946.

Furthauer, M. and Gonzalez-Gaitan, M. (2009). Endocytic regulation of notch signalling during development. Traffic 10, 792–802.

Gotz, M. and Huttner, W. B. (2005). The cell biology of neurogenesis. Nat Rev Mol Cell Biol 6, 777–88.

Gray, J. D., Kholmanskikh, S., Castaldo, B. S., Hansler, A., Chung, H., Klotz, B., Singh, S., Brown, A. M. and Ross, M. E. (2013). LRP6 exerts non-canonical effects on Wnt signaling during neural tube closure. Hum Mol Genet 22, 4267–81.

Herz, J., Goldstein, J. L., Strickland, D. K., Ho, Y. K. and Brown, M. S. (1991). 39-kDa protein modulates binding of ligands to low density lipoprotein receptor-related protein/alpha 2-macroglobulin receptor. J Biol Chem 266, 21232–8.

Hyvola, N., Diao, A., McKenzie, E., Skippen, A., Cockcroft, S. and Lowe, M. (2006). Membrane targeting and activation of the Lowe syndrome protein OCRL1 by rab GTPases. EMBO J 25, 3750–61.

Jeong, M. H., Ho, S. M., Vuong, T. A., Jo, S. B., Liu, G., Aaronson, S. A., Leem, Y. E. and Kang, J. S. (2014). Cdo suppresses canonical Wnt signalling via interaction with Lrp6 thereby promoting neuronal differentiation. Nat Commun 5, 5455.

Johnson, J. M., Castle, J., Garrett-Engele, P., Kan, Z., Loerch, P. M., Armour, C. D., Santos, R., Schadt, E. E., Stoughton, R. and Shoemaker, D. D. (2003). Genome-wide survey of human alternative pre-mRNA splicing with exon junction microarrays. Science 302, 2141–4.

Kettleborough, R. N., Busch-Nentwich, E. M., Harvey, S. A., Dooley, C. M., de Bruijn, E., van Eeden, F., Sealy, I., White, R. J., Herd, C., Nijman, I. J. et al. (2013). A systematic genome-wide analysis of zebrafish protein-coding gene function. Nature 496, 494–7.

Kimmel, C. B., Ballard, W. W., Kimmel, S. R., Ullmann, B. and Schilling, T. F. (1995). Stages of embryonic development of the zebrafish. Dev Dyn 203, 253–310.

Kounnas, M. Z., Argraves, W. S. and Strickland, D. K. (1992). The 39-kDa receptor-associated protein interacts with two members of the low density lipoprotein receptor family, alpha 2-macroglobulin receptor and glycoprotein 330. J Biol Chem 267, 21162–6.

Kressmann, S., Campos, C., Castanon, I., Furthauer, M. and Gonzalez-Gaitan, M. (2015). Directional Notch trafficking in Sara endosomes during asymmetric cell division in the spinal cord. Nat Cell Biol 17, 333–9.

Kur, E., Christa, A., Veth, K. N., Gajera, C. R., Andrade-Navarro, M. A., Zhang, J., Willer, J. R., Gregg, R. G., Abdelilah-Seyfried, S., Bachmann, S. et al. (2011). Loss of Lrp2 in zebrafish disrupts pronephric tubular clearance but not forebrain development. Dev Dyn 240, 1567–77.

Lee, H. O. and Norden, C. (2013). Mechanisms controlling arrangements and movements of nuclei in pseudostratified epithelia. Trends Cell Biol 23, 141–50.

Link, V., Shevchenko, A. and Heisenberg, C. P. (2006). Proteomics of early zebrafish embryos. BMC Dev Biol 6, 1.

Liu, T. L., Upadhyayula, S., Milkie, D. E., Singh, V., Wang, K., Swinburne, I. A., Mosaliganti, K. R., Collins, Z. M., Hiscock, T. W., Shea, J. et al. (2018). Observing the cell in its native state: Imaging subcellular dynamics in multicellular organisms. Science 360.

Lowery, L. A. and Sive, H. (2005). Initial formation of zebrafish brain ventricles occurs independently of circulation and requires the nagie oko and snakehead/atp1a1a.1 gene products. Development 132, 2057–67.

Luque, J. M., Morante-Oria, J. and Fairen, A. (2003). Localization of ApoER2, VLDLR and Dab1 in radial glia: groundwork for a new model of reelin action during cortical development. Brain Res Dev Brain Res 140, 195–203.

Marwaha, R. and Sharma, M. (2017). DQ-Red BSA Trafficking Assay in Cultured Cells to Assess Cargo Delivery to Lysosomes. Bio Protoc 7.

McCarthy, R. A., Barth, J. L., Chintalapudi, M. R., Knaak, C. and Argraves, W. S. (2002). Megalin functions as an endocytic sonic hedgehog receptor. J Biol Chem 277, 25660–7.

Mehta, Z. B., Pietka, G. and Lowe, M. (2014). The cellular and physiological functions of the Lowe syndrome protein OCRL1. Traffic 15, 471–87.

Nagai, M., Meerloo, T., Takeda, T. and Farquhar, M. G. (2003). The adaptor protein ARH escorts megalin to and through endosomes. Mol Biol Cell 14, 4984–96.

Nandez, R., Balkin, D. M., Messa, M., Liang, L., Paradise, S., Czapla, H., Hein, M. Y., Duncan, J. S., Mann, M. and De Camilli, P. (2014). A role of OCRL in clathrin-coated pit dynamics and uncoating revealed by studies of Lowe syndrome cells. Elife 3, e02975.

Nerli, E., Rocha-Martins, M. and Norden, C. (2020). Asymmetric neurogenic commitment of retinal progenitors involves Notch through the endocytic pathway. Elife 9.

Noakes, C. J., Lee, G. and Lowe, M. (2011). The PH domain proteins IPIP27A and B link OCRL1 to receptor recycling in the endocytic pathway. Mol Biol Cell 22, 606–23.

Norden, C. (2017). Pseudostratified epithelia - cell biology, diversity and roles in organ formation at a glance. J Cell Sci 130, 1859–1863.

Oltrabella, F., Jackson-Crawford, A., Yan, G., Rixham, S., Starborg, T. and Lowe, M. (2021). IPIP27A cooperates with OCRL to support endocytic traffic in the zebrafish pronephric tubule. Hum Mol Genet.

Oltrabella, F., Pietka, G., Ramirez, I. B., Mironov, A., Starborg, T., Drummond, I. A., Hinchliffe, K. A. and Lowe, M. (2015). The Lowe syndrome protein OCRL1 is required for endocytosis in the zebrafish pronephric tubule. PLoS Genet 11, e1005058.

Parisi, M. A. (2009). Clinical and molecular features of Joubert syndrome and related disorders. Am J Med Genet C Semin Med Genet 151C, 326–40.

Perez Bay, A. E., Schreiner, R., Benedicto, I., Paz Marzolo, M., Banfelder, J., Weinstein, A. M. and Rodriguez-Boulan, E. J. (2016). The fast-recycling receptor Megalin defines the apical recycling pathway of epithelial cells. Nat Commun 7, 11550.

Ramirez, I. B., Pietka, G., Jones, D. R., Divecha, N., Alia, A., Baraban, S. C., Hurlstone, A. F. and Lowe, M. (2012). Impaired neural development in a zebrafish model for Lowe syndrome. Hum Mol Genet 21, 1744–59.

Ro, H., Soun, K., Kim, E. J. and Rhee, M. (2004). Novel vector systems optimized for injecting in vitro-synthesized mRNA into zebrafish embryos. Mol Cells 17, 373–6.

Schindelin, J., Arganda-Carreras, I., Frise, E., Kaynig, V., Longair, M., Pietzsch, T., Preibisch, S., Rueden, C., Saalfeld, S., Schmid, B. et al. (2012). Fiji: an open-source platform for biological-image analysis. Nat Methods 9, 676–82.

Shah, M., Baterina, O. Y., Jr., Taupin, V. and Farquhar, M. G. (2013). ARH directs megalin to the endocytic recycling compartment to regulate its proteolysis and gene expression. J Cell Biol 202, 113–27.

Sheen, V. L., Ganesh, V. S., Topcu, M., Sebire, G., Bodell, A., Hill, R. S., Grant, P. E., Shugart, Y. Y., Imitola, J., Khoury, S. J. et al. (2004). Mutations in ARFGEF2 implicate vesicle trafficking in neural progenitor proliferation and migration in the human cerebral cortex. Nat Genet 36, 69–76.

Spoelgen, R., Hammes, A., Anzenberger, U., Zechner, D., Andersen, O. M., Jerchow, B. and Willnow, T. E. (2005). LRP2/megalin is required for patterning of the ventral telencephalon. Development 132, 405–14.

Takita, S. and Seko, Y. (2020). eys (+/-) ; lrp5 (+/-) Zebrafish Reveals Lrp5 Can Be the Receptor of Retinol in the Visual Cycle. iScience 23, 101762.

Taverna, E., Gotz, M. and Huttner, W. B. (2014). The cell biology of neurogenesis: toward an understanding of the development and evolution of the neocortex. Annu Rev Cell Dev Biol 30, 465–502.

Vacaru, A. M., Unlu, G., Spitzner, M., Mione, M., Knapik, E. W. and Sadler, K. C. (2014). In vivo cell biology in zebrafish - providing insights into vertebrate development and disease. J Cell Sci 127, 485–95.

van Rahden, V. A., Brand, K., Najm, J., Heeren, J., Pfeffer, S. R., Braulke, T. and Kutsche, K. (2012). The 5-phosphatase OCRL mediates retrograde transport of the mannose 6-phosphate receptor by regulating a Rac1-cofilin signalling module. Hum Mol Genet 21, 5019–38.

Veth, K. N., Willer, J. R., Collery, R. F., Gray, M. P., Willer, G. B., Wagner, D. S., Mullins, M. C., Udvadia, A. J., Smith, R. S., John, S. W. et al. (2011). Mutations in zebrafish lrp2 result in adult-onset ocular pathogenesis that models myopia and other risk factors for glaucoma. PLoS Genet 7, e1001310.

Vicinanza, M., Di Campli, A., Polishchuk, E., Santoro, M., Di Tullio, G., Godi, A., Levtchenko, E., De Leo, M. G., Polishchuk, R., Sandoval, L. et al. (2011). OCRL controls trafficking through early endosomes via PtdIns4,5P(2)-dependent regulation of endosomal actin. EMBO J 30, 4970–85.

Wang, B., He, W., Prosseda, P. P., Li, L., Kowal, T. J., Alvarado, J. A., Wang, Q., Hu, Y. and Sun, Y. (2021). OCRL regulates lysosome positioning and mTORC1 activity through SSX2IP-mediated microtubule anchoring. EMBO Rep 22, e52173.

Wilkinson, R. N., Elworthy, S., Ingham, P. W. and van Eeden, F. J. (2013). A method for high-throughput PCR-based genotyping of larval zebrafish tail biopsies. Biotechniques 55, 314–6.

Willnow, T. E. and Christ, A. (2017). Endocytic receptor LRP2/megalin-of holoprosencephaly and renal Fanconi syndrome. Pflugers Arch 469, 907–916.

Willnow, T. E., Goldstein, J. L., Orth, K., Brown, M. S. and Herz, J. (1992). Low density lipoprotein receptor-related protein and gp330 bind similar ligands, including plasminogen activator-inhibitor complexes and lactoferrin, an inhibitor of chylomicron remnant clearance. J Biol Chem 267, 26172–80.

Yamamoto, H., Umeda, D., Matsumoto, S. and Kikuchi, A. (2017). LDL switches the LRP6 internalization route from flotillin dependent to clathrin dependent in hepatic cells. J Cell Sci 130, 3542–3556.

Zhang, X., Hartz, P. A., Philip, E., Racusen, L. C. and Majerus, P. W. (1998). Cell lines from kidney proximal tubules of a patient with Lowe syndrome lack OCRL inositol polyphosphate 5-phosphatase and accumulate phosphatidylinositol 4,5-bisphosphate. J Biol Chem 273, 1574–82.

Zhao, X., Garcia, J. Q., Tong, K., Chen, X., Yang, B., Li, Q., Dai, Z., Shi, X., Seiple, I. B., Huang, B. et al. (2021). Polarized endosome dynamics engage cytoplasmic Par-3 that recruits dynein during asymmetric cell division. Sci Adv 7.

